# Estimation of narrowband amplitude and phase from electrophysiology signals for phase-amplitude coupling studies: a comparison of methods

**DOI:** 10.1101/392886

**Authors:** Juan L.P. Soto, Felipe V.D. Prado, Etienne Combrisson, Karim Jerbi

## Abstract

Many functional connectivity studies based on electrophysiological measurements, such as electro- and magnetoencephalography (EEG/MEG), start their investigations by extracting a narrowband representation of brain activity time series, and then computing their envelope amplitudes and instantaneous phases, which serve as inputs to subsequent data processing. The two most popular approaches for obtaining these narrowband amplitudes and phases are: bandpass filtering followed by Hilbert transform (we call this the Hilbert approach); and convolution with wavelet kernels (the wavelet approach). In this work, we investigate how these two approaches perform in detecting the phenomenon of phase-amplitude coupling (PAC), whereby the amplitude of a high-frequency signal is driven by the phase of a low-frequency signal. The comparison of both approaches is carried out by means of simulated brain activity, from which we run receiver operating characteristic (ROC) analyses, and of experimental MEG data from a visuomotor coordination study. The ROC analyses show that both approaches have comparable accuracy, except in the presence of interfering signals with frequencies near the high-frequency band. As for the visuomotor data, the most noticeable impact of the choice of approach was observed when evaluating task-based changes in PAC between the delta (2-5 Hz) and the high-gamma (60-90 Hz) frequency bands, as we were able to identify widespread brain areas with statistically significant effects only with the Hilbert approach. These results provide preliminary evidence of the advantages of the Hilbert approach over the wavelet approach, at least in the context of PAC estimates.

## 1 Introduction

In Neuroscience, one of the most dynamic and fastest-evolving fields of study nowadays is functional connectivity, i.e. how two or more sources of neuronal activity coordinate in response to a given experimental task. Electro- and magnetoencephalography (EEG/MEG) are two imaging modalities particularly suitable for connectivity studies due to their high temporal resolution (on the order of milliseconds), allowing the analysis of interaction patterns that vary over time and frequency (Baillet et al., 2001; Darvas and Leahy, 2007; Schoffelen and Gross, 2009; Gross et al., 2013). One of the most promising applications of EEG/MEG to functional interaction analysis is the estimation of phase-amplitude coupling (PAC, also called nested oscillations), a phenomenon whereby the envelope amplitude of a high-frequency (HF) signal is driven by the instantaneous phase of a low-frequency (LF) one (Canolty et al., 2006; Jensen and Colgin, 2007; Canolty and Knight, 2010; Aru et al., 2015; Nakhnikian et al., 2016); the HF and LF signals typically come from the neuronal activity of a single location, but they may also refer to separate areas (Maris et al., 2011; van der Meij et al., 2012; Guirgis et al., 2015).

Most studies dealing with PAC apply one of the following time-frequency approaches to extract narrowband representations of brain activity and to calculate their envelope amplitude and instantaneous phase: band-pass filtering of the activity signal followed by Hilbert transform (we will call this the *Hilbert approach*) (Cohen, 2008; Kramer et al., 2008; Voytek et al., 2010, 2013; McGinn and Valiante, 2014; Voytek et al., 2015a,b; Niknazar et al., 2015; Ninomiya et al., 2015; Smith et al., 2015; Xu et al., 2015; Blain-Moraes et al., 2015; van Wijk et al., 2015); or convolution with a continuous wavelet kernel (we will call this the *wavelet approach*), sometimes preceded by band-pass filtering (Maris et al., 2011; Canolty et al., 2012; Florez et al., 2015; McGinn and Valiante, 2014; Kajihara et al., 2015; O’Connell et al., 2015; Mizuhara et al., 2015; Sweeney-Reed et al., 2016; Vandenbroucke et al., 2015; van Driel et al., 2015). The two approaches are formally equivalent, since both of them can be formulated as linear convolutions with the activity time series, and one can make them to be identical with an adequate choice of parameters (particularly those of the band-pass filters) (Bruns, 2004; Kiebel et al., 2005). Moreover, studies using simulated and real electrophysiological data have demonstrated that there are only very small quantitative differences between the approaches when they are applied to the estimation of narrowband amplitude envelopes (Bruns, 2004) and to the computation of synchrony between the instantaneous phases at two separate regions (Le Van Quyen et al., 2001). These similarities in performance, however, do not take into account that, in practice, bandpass filters with frequency responses similar to those of wavelet convolutions have unattractive properties in terms of frequency discrimination, and therefore are seldom applied to connectivity studies. In the specific case of PAC estimations, another reason why the Hilbert and the wavelet approaches may not have comparable performance has to do with particular features of the signals employed in this type of analysis. The extraction of the amplitude envelope and the instantaneous phase from narrowband signals requires analytic time series (i.e. with no spectral components for negative frequencies), a condition satisfied by the Hilbert transform but not by the wavelet convolution; the latter yields only approximately analytic representations, and only if the central frequency of the wavelet kernel is sufficiently high (Quian Quiroga et al., 2002). Since the phases for the PAC computations are often computed for very low signal frequencies (as low as 2 Hz), the use of wavelets in this context may result in inaccurate assessments of coupling.

The main goal of the present work is evaluate quantitatively the performance of the Hilbert and the wavelet approaches in the extraction of the signal representations required for phase-amplitude coupling estimations from EEG/MEG data. We measure performance with ROC curves and synthetic activity time series with varying noise levels, durations and coupling strengths. We also illustrate the differences between the Hilbert and the wavelet techniques with real MEG data acquired during a visuomotor coordination study. By carrying out these tests, we intend to call attention to characteristics of these methods that must be dealt with carefully lest imprecise coupling measures be obtained, not only in PAC studies.

## 2 Methods

In figure 1 we present an overview of the main transformations we performed on brain activity time signals in order to obtain estimates of PAC. First, we extracted analytic representations of the components of the input signal lying within the frequency bands of interest (in the example of figure 1, 4-7 Hz and 30-60 Hz). With the Hilbert approach, these representations were obtained in two steps, i.e. band-pass filtering followed by Hilbert transforms, whereas with the wavelet approach, the convolution of the input with the wavelet kernels directly resulted in complex-valued, narrowband time series. With either approach, we then computed the instantaneous phase (for the lower band) and the envelope amplitude (for the higher band) time series. Both PAC metrics used here (as detailed in subsection 2.3) depend on the probability distributions of the amplitude envelopes as functions of the instantaneous phases – specifically, on how much the distributions differ from those of a uniform random variable.

**Figure 1:**
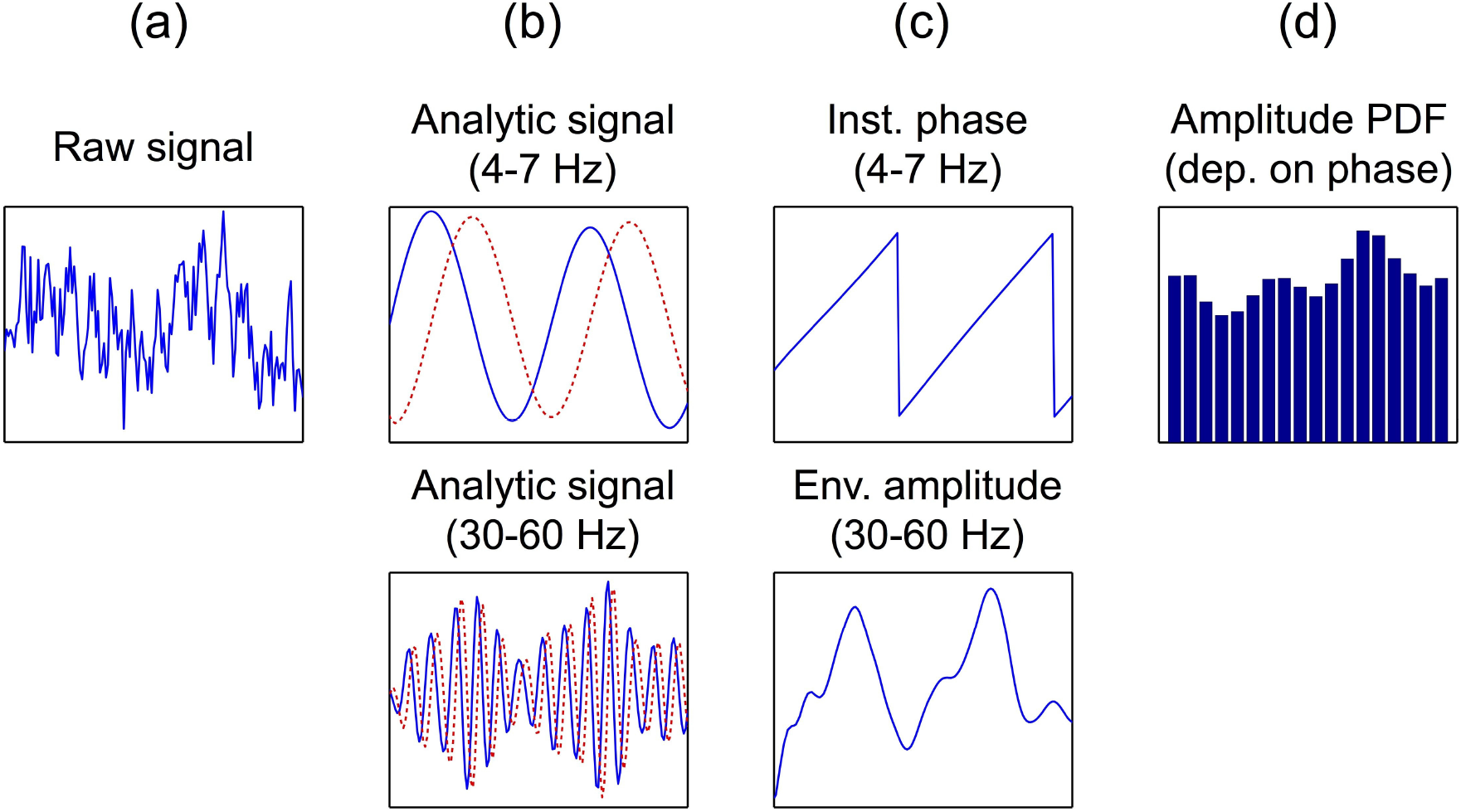
Main steps in our procedure to compute PAC from brain activity time series. (a) Input activity signal. (b) Analytic representations of the original signal’s components that lie within the theta (top plot) and low-gamma (bottom plot) frequency bands; with the Hilbert approach, these representations are obtained by band-pass filtering followed by applying the Hilbert transform, while with the wavelet approach, convolution with the wavelet kernels result directly in the analytic signals. (In these plots, blue lines represent the signals’ real parts, and red lines indicate their imaginary parts.) (c) Theta instantaneous phase (top plot) and low-gamma envelope amplitude (bottom plot) time series, computed from the analytic signals shown in (b). (d) Probability distribution of the low-gamma envelope amplitude values as function of the theta instantaneous phases, computed from the time series shown in (c). The PAC metrics we implemented in this work, discussed in subsection 2.3, are ways to assess how much this distribution departs from that of an uniform random variable.

### 2.1 The Hilbert approach

Let time series *x*(*t*) be a measure of localized brain activity. For instance, *x*(*t*) may be the recorded voltages (or magnetic fields) at a specific EEG (or MEG) sensor or, in source space analyses, the estimate of the electric current density at a chosen brain region reconstructed from EEG or MEG data. The first step in the Hilbert approach is to apply a band-pass filter to *x*(*t*) to obtain *x_B_*(*t*), a narrowband version of the original time series with components (ideally) restricted to frequency band *B* – throughout this work, the bands investigated were delta (*δ*, 2-5 Hz), theta (*θ*, 4-7 Hz), low-gamma (*γ*_*L*_, 30-60 Hz) and high-gamma (*γ*_*H*_, 60-90 Hz). All filters in the present study had a number of coefficients equal to *N*_time_/3 or its nearest integer, *N*_time_ being the number of time points in *x*(*t*), and they were computed with an algorithm that seeks to minimize the least-squares error between an ideal bandpass filter and the actual one (function `firls.m` in Matlab). The coefficients were then weighted by a Kaiser window with parameter 3, and finally the filtering was performed in the forward and backward directions to prevent phase distortions (function filtfilt.m in Matlab). For a given frequency band B = [*f*_1_, *f*_2_], the desired passband used in the filter design algorithms was [*f*_1_ — 0.1△*f*, *f*_2_ + 0.1△*f*] (where Δ*f* = *f*_2_ – *f*_1_), to ensure that the full band of interest passes through the filter. The narrowband signal is then convolved with a function whose frequency response is the Heaviside step function, resulting in *x_H_*(*t*) = Re[*x_H_*(*t*)] + *j*{Im[*x_H_*(*t*)]}, the analytic representation of *x*_*B*_(*t*) (throughout this work, *j* = 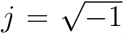). The expressions to compute the Hilbert-based narrowband envelope amplitude and instantaneous phase are, respectively:

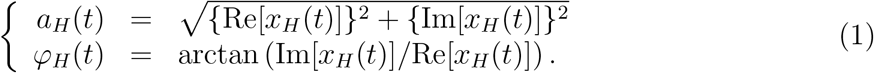

### 2.2 The wavelet approach

In this approach, we obtain *x_ν_*(*t*), a complex, narrowband representation of the original time series *x*(*t*), by convolving the latter with a continuous Morlet wavelet kernel, a Gaussian-weighted, complex sinusoidal function whose expression is (Pantazis et al., 2009):

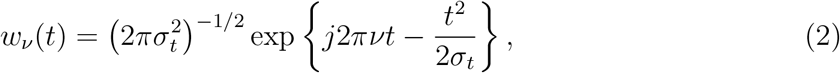

where *ν* is the kernel s central frequency. In the time domain, <*w_ν_*(*t*)< has the shape of a centered Gaussian function with standard deviation *σ_t_*, and in the frequency domain this kernel is also Gaussian-shaped, with peak at *ν* and standard deviation *σ_ν_* = 1/(2*πσ_t_*); we can also express these standard deviations in terms of the number of cycles *N*_cyc_, or the number of wavelet cycles within a 6*σ_t_* time interval, which is given by *N*_cyc_ = 2*πσ_t_ν = ν/σ_ν_*. Due to the frequency response properties of the Morlet wavelets, the use of a single kernel to identify signal components lying within the frequency band of interest may lead either to strong attenuation of components far from the center of the band, or to the detection of components lying outside the desired range. The selection of the optimal values for *N*_cyc_ and *N*_ker_ (the number of wavelet kernels with central frequencies uniformly spaced between *f*_1_ and *f*_2_) was performed with simulated current density time series, as will be detailed below. Similarly to equation (1), the wavelet-based envelope amplitude *α_W,ν_*(*t*) and the instantaneous phase *φ_w,v_*(*t*) can then be expressed for each *ν* in terms of the real and imaginary parts of *x_ν_*(*t*):

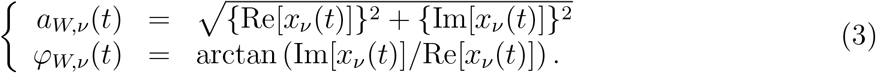

Figure 2 displays a comparison between both approaches in terms of frequency response, when they are set to discriminate signal components within the *δ* and the *γ*_*H*_ frequency bands. To generate these plots, we created signals *x*_0_(*t*) = cos(2*πf*_0_*t*) with length 1 second, sampling frequency 1 kHz and several values of *f*_0_, then computed the average envelope amplitude of the output signal when both approaches are applied to *x*_0_(*t*).

**Figure 2:**
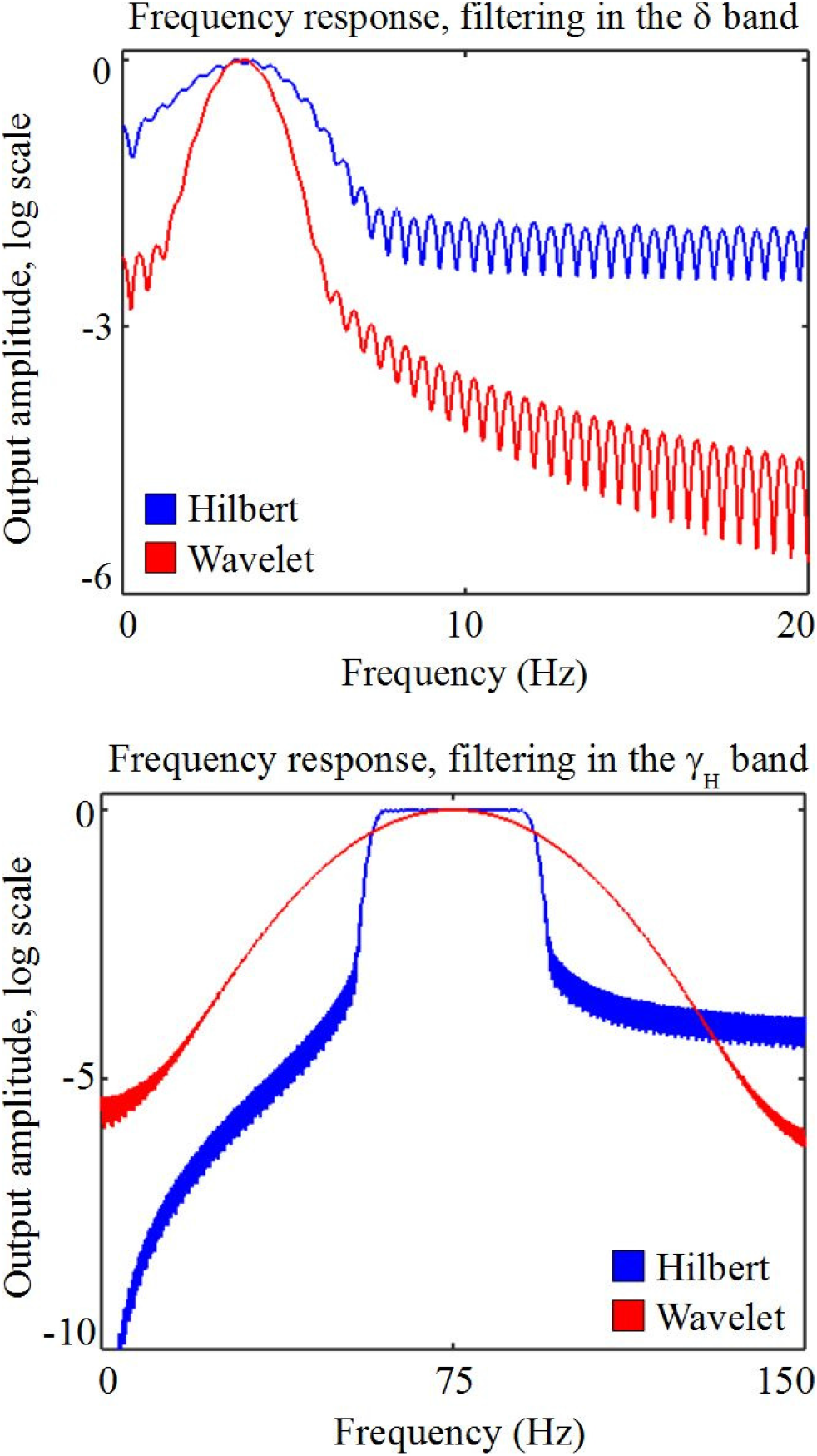
Frequency response of the Hilbert (blue line) and the wavelet (red line) approaches when they are set to detect signals in the *δ* (top plot) and *γ*_*H*_ (bottom plot) frequency bands. These plots display, as a function of frequency *f*_0_ and on a logarithmic scale, the average output amplitude when both approaches are applied to signal *x*_0_ = cos(2*πf*_0_*t*).

### 2.3 Phase-amplitude coupling

The phenomenon of phase-amplitude coupling occurs when the envelope amplitude of a fast narrowband time series oscillates in synchrony with a slower signal (Canolty et al., 2006; Jensen and Colgin, 2007). Several measures have been proposed in the literature to assess the amount of PAC between two oscillations; in order to prevent that a poor performance be due merely to the choice of PAC measure, we employ two such metrics in the present study.

One is based on the Kullback-Leibler divergence (Tort et al., 2010), which has been demonstrated to provide better performance in terms of detection of strong effects and rejection of false positives (Soto and Jerbi, 2012). This PAC estimator seeks to quantify how much the distribution of the high-frequency amplitudes *a*_hi_(*t*) as a function of the low-frequency phases *φ*_lo_(*t*) departs from an uniform distribution, the latter indicating no relationship between *a*_hi_(*t*) and *φ*_lo_(*t*). Specifically, we compute the following:

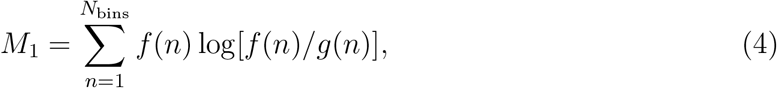

where *f*(*n*) is the (discrete) empirical distribution of *a*_hi_(*t*), *g*(*n*) is the uniform distribution, and *N*_bins_ = 18 is the number of bins in which the *φ*_lo_(*t*) values are divided in order to estimate *f*(*n*) (i.e. the number of points in either distribution).

The other PAC estimator used here measures the linear relationship between *a*_hi_(*t*) and exp{*jφ*_lo_(*t*)}, normalized by the mean energy of the high-frequency amplitude (Ozkurt, 2012):

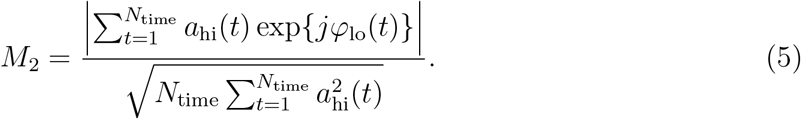

Unlike *M*_1_, the calculation of *M*_2_ does not require explicitly the estimation of *f*(*n*), but it is related to the statistical dependence between the phases and the amplitudes, since *M*_2_ is very small when *f*(*n*) is near-uniform.

With the Hilbert approach, the low-frequency band provides one time series *φ*_lo_(*t*), and the high-frequency band, one time series *a*_hi_(*t*); from these two time series a single PAC estimate (*M*_1_ or *M*_2_) is computed. On the other hand, if more than one wavelet kernel is used for either frequency band (say, *N*_ker_ = 3 for both bands), then there will be multiple PAC estimates from these signals (3^2^ = 9 estimates for this example). For our comparisons with the wavelet approach, the final PAC statistic was the mean over these multiple initial PAC estimates.

### 2.4 ROC analysis with simulated data

The synthetic activity signals used in our comparisons between the Hilbert and the wavelet approach consisted of a summation of three sinusoidal waves and noise. Two of the sinusoids had frequencies falling within intervals *B*_hi_ = [*f*_hi,1_, *f*_hi,2_] and *B*_lo_ = [*f*_lo,1_, *f*_lo,2_], respectively, and the envelope amplitude of the faster sinusoid was coupled with the slower one; the other sinusoid oscillated at a frequency outside either *B*_lo_ or *B*_hi_, and was uncorrelated with the other two sinusoids. The mathematical expression of the simulated activity signals, adapted from the expression proposed by Tort et al. (2010), is given by:

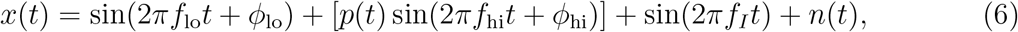

where *n*(*t*) was zero-mean Gaussian noise with variance σ^2^, *φ*_lo_ and *φ*_hi_ were randomly sampled from a uniform distribution between 0 and 2*π*, *f*_lo_ and *f*_hi_ were randomly sampled from a uniform distribution on the intervals *B*_lo_ and *B*_hi_, respectively, f_I_ was a constant, and time series *p*(*t*) had the expression:

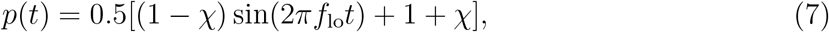

*χ* being the coupling parameter. If *χ* = 1, there was no coupling between the *f*_hi_ and the *f*_lo_ sinusoids, while values of *χ* close to 0 represented strong coupling. The inclusion of the term sin(2*πf*/1) in the expression of *x*(*t*) was meant to investigate the impact of signal components not within *B*_lo_ and *B*_hi_ in the estimation of PAC, since brain activity is usually not restricted to the two frequency bands under study.

The generation of the synthetic time series *x*(*t*) and the subsequent PAC computation were dependent on the following parameters:

- sampling frequency *f_s_*;
- signal length *T* = *N*_time_/*f_s_*;
- noise variance *σ*^2^;
- low- and high-frequency bands *B*_lo_ (*δ* or *θ*) and *B*_hi_ (*γ*_*L*_ or *γ*_*H*_);
- coupling parameter *χ*;
- PAC estimator, i.e. *M*_1_ or *M*_2_ (equations (4) and (5));
- frequency *f_I_* of signal component outside *B*_lo_ and *B*_hi_.

Our first goal was to determine the values of *N_cyc_* (the number of cycles in the wavelet kernels) and *N_ker_* (the number of kernels spanning the frequency bands of interest) that provided the best classification performance. For this, we chose a fixed parameter set (*T* = 1s, *f_s_* = 500Hz, *χ* = 0.5, *σ* = 1, PAC estimator *M*_1_) and, for each combination of *B*_lo_ and *B*_hi_, varied first *N*_cyc_ with *N*_ker_ = 1, then *N*_ker_ with the optimal *N*_cyc_. Having chosen the wavelet parameters, we then proceeded to the comparison of approaches, changing the values for the time series parameters (*χ, σ* etc.).

The ROC analysis for a given choice of parameter values was performed by creating 10000 realizations of *x*(*t*), 5000 with *χ* = 1 and 5000 with a specified value of *χ* < 1. From each realization, we extracted one high-frequency amplitude envelope time series *a*_hi_(*t*) and one low-frequency instantaneous phase time series *φ*_lo_(*t*) with the Hilbert transform; with the wavelet approach, depending on the choice of number of kernels, we obtained a single or several pairs of phase and amplitude time series, as discussed in subsection 2.2. PAC was then estimated with one of the two methods described in subsection 2.3 – if *N*_ker_ > 1 with the wavelet approach, we took the mean over all PAC estimates computed from every combination of amplitude and phase time series. Finally, the ROC curve could be determined by applying a variable threshold *ϑ* to the 10000 computed values of the chosen PAC metric: for a given threshold value, the true positive fraction (TPF) was given by the proportion of simulated *x*(*t*) with *χ* < 1 that resulted in a PAC estimate larger than or equal to *ϑ*, while the proportion of the *x*(*t*) with *χ* = 1 with above-threshold PAC yielded the false positive fraction (FPF).

### 2.5 MEG experimental data

To illustrate the differences between the two approaches when estimating PAC based on real electrophysiology signals, we ran analyses on MEG data acquired from 15 subjects during a visuomotor coordination study; details about the experimental paradigm and data preprocessing can be found elsewhere (Jerbi et al., 2004, 2007). Briefly, the experiment consisted of two conditions: visuomotor coordination (VM), in which the subjects were required to watch a rotating cube on a screen in front of them and simultaneously prevent the rotations by operating a trackball with their right hand; and rest (R), in which the subjects stared at a still cube and did not perform any other activity. Steady-state data was recorded during 8-minute sessions, in which every 8 to 12 seconds the conditions were switched. The 8-12 s data segments were visually inspected for eye blinks and other movement artifacts; those that were not discarded were subjected to template-matching to deal with heart-beat effects, and then divided into 1-second trials. The data sampling frequency was 312.5 Hz.

Each trial from each subject *m, m* ∈ {1,…, 15}, was converted into an estimated cortical current density map by means of a regularized minimum-norm inverse method (Tikhonov and Arsenin, 1977; Okada, 2003), which in turn was based on a simplified spherical head model (Wolters et al., 2006; Mosher et al., 1999); these procedures we implemented with BrainStorm, an open source software package (Tadel et al., 2011). The current density map from trial *l* of condition *k* (*k* ∈ {VM, R}) can be written as a matrix 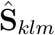, with dimension *N*_sources_ × *N*_time_. From each 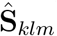 computed two cortical maps 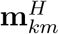 and 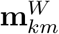, each with dimension *N*_sources_ × 1, of the PAC estimate *M* (one map with the Hilbert approach, and one with the wavelet approach). In order to test whether PAC (say, with the Hilbert approach) changes significantly from R to VM at a given spatial location s for subject m, we employed the statistic

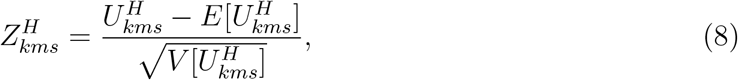

where 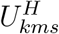 is the Wilcoxon rank sum test statistic based on the *s*-th element of all column vectors 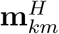, with mean 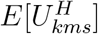 and variance 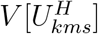. The standardization in equation (8) results in a statistic 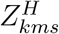 which is approximately normal (Hollander et al., 2013), and by doing this we intended to compensate for different trial numbers across subjects. Repeating this procedure for all subjects and both conditions yielded 30 values of 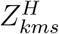 corresponding to location *s*, 15 per condition. From these 30 values we then computed a t-statistic 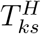, whose probability distribution was estimated empirically with a nonparametric resampling method (Nichols and Holmes, 2001). The resampling consisted in choosing randomly a sample of the 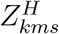, multiplying them by −1, and then computing the surrogate 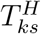 with the new values of 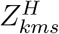; repeating this process several times provided the samples to the empirical distribution. Given the large number of statistical tests that must be performed (one for each *s*), we used these distributions to correct for multiple comparisons, based on a procedure that controls for the false discovery rate (FDR), or the expected number of false positives (Benjamini and Hochberg, 1995; Nichols and Hayasaka, 2003).

## 3 Results

### 3.1 Parameter selection

Table 1 lists the values of the areas under the ROC curves (AUCs) computed with simulated time series and the wavelet approach, with different values of the number of wavelet cycles *N*_cyc_ and for each combination of low-frequency (*δ* and *θ*) and high-frequency (*γ*_*L*_ and *γ*_*H*_) pairs. For this choice of parameters (as described in subsection 2.4), it was found that 4 wavelet cycles provided the best classification performance when the high-frequency band was *γ*_*H*_, while *N*_cyc_ = 3 was the optimal number of cycles in PAC estimations with *γ*_*L*_ as the high-frequency band. Next, we performed the same analysis with simulations, but with the optimal values of *N*_cyc_ and with different values of the number of wavelet kernels *N*_ker_; the resulting AUCs appear in table 2. It can be seen that more wavelet kernels led to an improvement for the wavelet approach, except for the frequency band pair *θ* — *γ*_*L*_; however, given that over all band pairs the change in performance was smaller than 2.5%, and that higher *N*_ker_ necessarily result in longer processing times, we chose *N*_ker_ = 1 for our subsequent comparisons between the approaches.

**Table 1:** Values of areas under ROC curves (AUCs) as functions of the number of cycles *N*_cyc_ of the wavelet kernels for each pair of frequency band of interest. The parameters chosen for the simulated time series used in these computations were: *T* = 1s, *f_s_* = 500Hz, *χ* = 0.5, *σ* = 1, PAC method *M*_1_ and *N*_ker_ = 1. Optimal values for each frequency band pair are highlighted.

| | Number of cycles $N_{\text{cyc}}$ | | | | | |
| --- | --- | --- | --- | --- | --- | --- |
|  | 1 | 2 | 3 | 4 | 5 | 6 |
| $\delta\text{-}\gamma_H$ | .7330 | .8435 | .8958 | **.9005** | .8934 | .8740 |
| $\delta\text{-}\gamma_L$ | .7815 | .8894 | **.9091** | .8699 | .8208 | .7583 |
| $\theta\text{-}\gamma_H$ | .7534 | .8625 | .9064 | **.9189** | .9076 | .9006 |
| $\theta\text{-}\gamma_L$ | .7764 | .9081 | **.9138** | .8869 | .8376 | .7926 |

**Table 2:** Values of areas under ROC curves (AUCs) as functions of the number of wavelet kernels *N*_ker_ for each pair of frequency band of interest. The parameters chosen for the simulated time series used in these computations were: *T* = 1s, *f_s_* = 500Hz, *χ* = 0.5, *σ* =1, PAC method *M*_1_ and the values of Ncyc highlighted in table 1. Optimal values for each frequency band pair are highlighted.

| | Number of kernels $N_{\text{ker}}$ | | |
| --- | --- | --- | --- |
|  | 1 | 3 | 5 |
| $\delta\text{--}\gamma_H$ | .9005 | .9150 | **<u>.9266</u>** |
| $\delta\text{--}\gamma_L$ | .9091 | .8962 | **<u>.9122</u>** |
| $\theta\text{--}\gamma_H$ | .9189 | .9199 | **<u>.9229</u>** |
| $\theta\text{--}\gamma_L$ | **<u>.9138</u>** | .9050 | .9066 |

### 3.2 ROC analysis

The areas of the ROC curves (AUCs) obtained from the Hilbert and wavelet approaches for different values of the parameters listed in section 2.4 are presented in figures 3-8; in each of these tables, the AUC values are presented as a function of a given parameter (or of two parameters, as in figure 7), while the other parameters are kept constant. As figures 3 through 7 show (with a single exception), there were only very small differences in classification accuracy between the Hilbert and the wavelet approaches when we varied the coupling strength, the noise amplitude, the data length (either increasing time or sampling frequency) and the interacting frequency bands. The exception appears in the lower plot of figure 7, for which we changed the method for estimating PAC – i.e. the advantage of using the wavelet approach was slightly higher than in the other figures mentioned above; still, for both approaches, the use of *M*_2_ (equation (5)) instead of *M*_1_ (4) to compute PAC resulted in lower AUC values, which is in line with findings we presented elsewhere (Soto and Jerbi, 2012) regarding the benefits of choosing the Kullback-Leibler divergence-based expression over other techniques to assess cross-frequency coupling. On the other hand, the presence of a single sinusoidal function added to the simulated time series, as presented in figure 8, caused a noticeable worsening in the accuracy of the wavelet approach in comparison with the Hilbert approach, especially for sinusoid frequencies near the lower limit of the *γ*_*H*_ band (60 Hz) – this decrease in AUC values was not observed for *f_I_* values near the upper limit of the *δ* band (5 Hz), thus they do not appear in figure 8. From these plots we can also observe that this degradation linked to the wavelet approach occurred regardless of whether the sinusoidal interference was exclusive to a single condition or not, even though the disparities in performance were not as strong in the case when *f_I_* = 0 only during the task condition (i.e. in signals with *χ* < 1).

**Figure 3:**
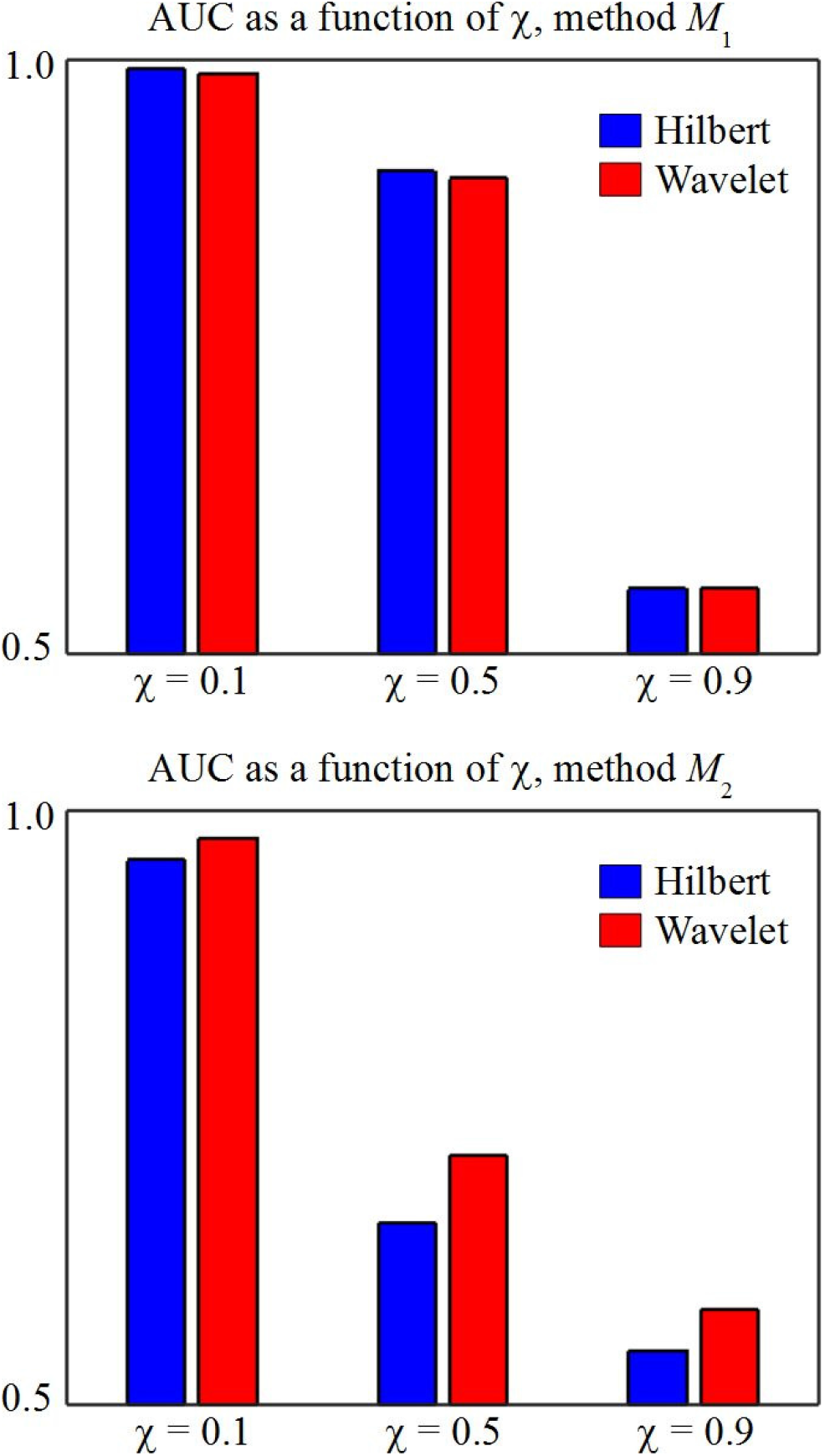
Areas under ROC curves for different values of the coupling parameter *χ*; blue bars are used for the Hilbert approach, and red bars for the wavelet approach. The PAC method was *M*_1_ for the top plot, and *M*_2_ for the bottom plot. The values for the other parameters are: *T* = 1 s, *σ* = 1, *B*_lo_ = *δ*, *B*_hi_ = *γ*_*H*_, *f_s_* = 500 Hz and *f_I_* = 0.

**Figure 4:**
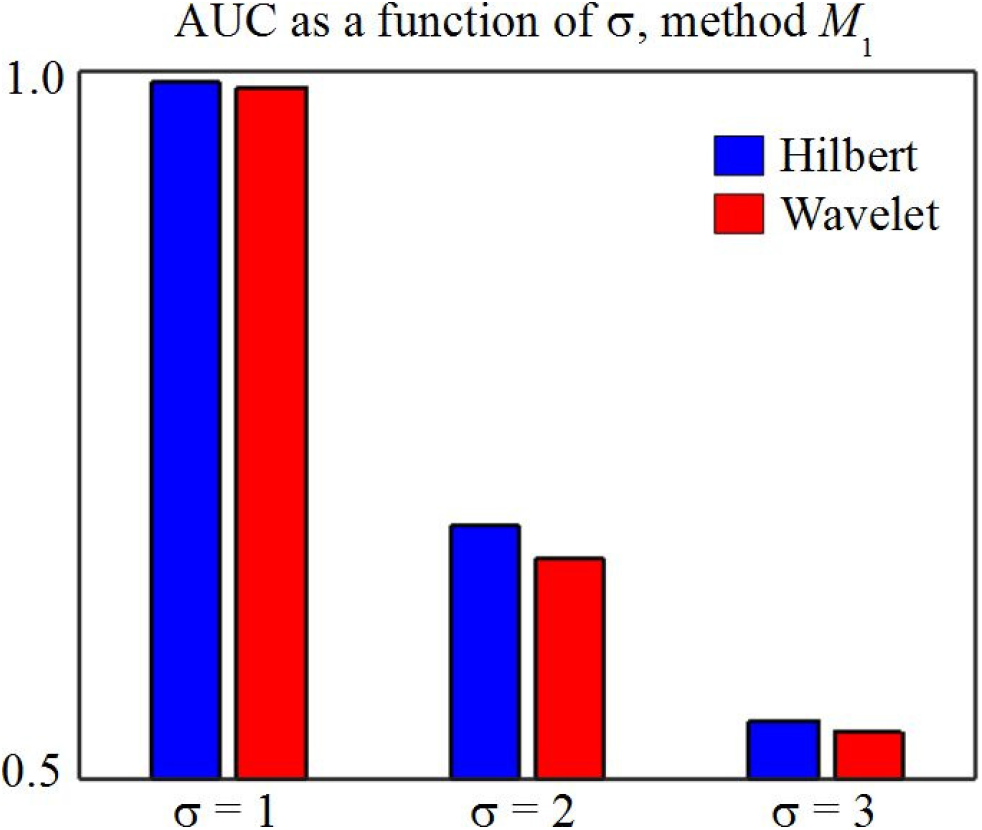
Areas under ROC curves for different values of the noise standard deviation *σ*; blue bars are used for the Hilbert approach, and red bars for the wavelet approach. The values for the other parameters are: *T* = 1 s, *χ* = 0.1, *B*_lo_ = *δ*, *B*_hi_ = *γ*_*H*_, *f_s_* = 500 Hz, *f_I_* = 0 and PAC method *M*_1_.

**Figure 5:**
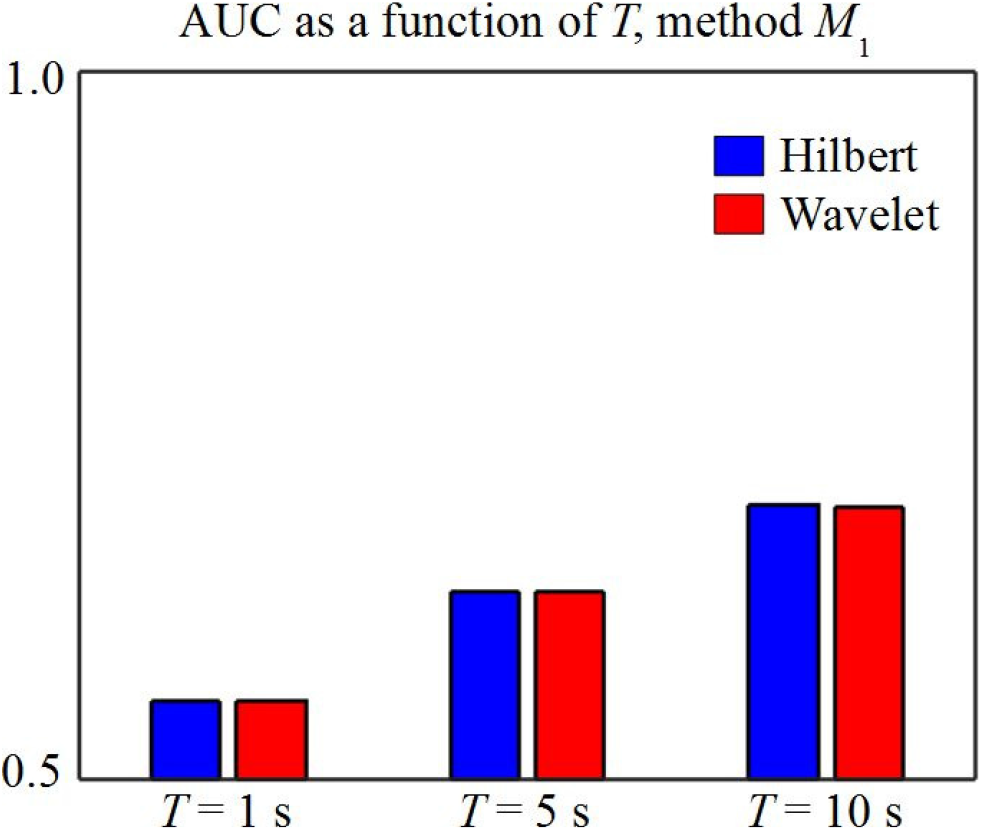
Areas under ROC curves for different values of the signal length *T* in seconds; blue bars are used for the Hilbert approach, and red bars for the wavelet approach. The values for the other parameters are: *χ* = 0.5, *σ* = 1, *B*_lo_ = *δ*, *B*_hi_ =, *f_s_* = 500 Hz, *f_I_* = 0 and PAC method *M*_1_.

**Figure 6:**
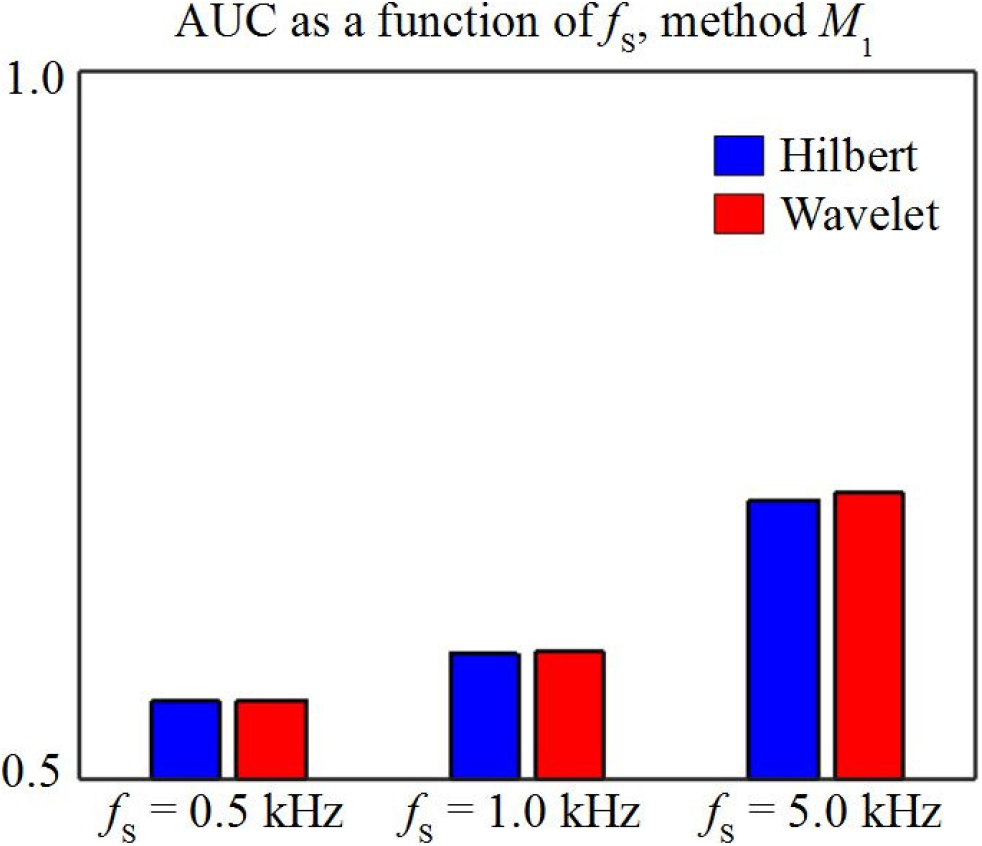
Areas under ROC curves for different values of the sampling frequency *f_s_* in kHz; blue bars are used for the Hilbert approach, and red bars for the wavelet approach. The values for the other parameters are: *χ* = 0.5, *T* = 1 s, *σ* =1, *B*_lo_ = *δ*, *B*_hi_ = *γ*_*H*_, *f_s_* = 500 Hz, *f_I_* = 0 and PAC method *M*_1_.

**Figure 7:**
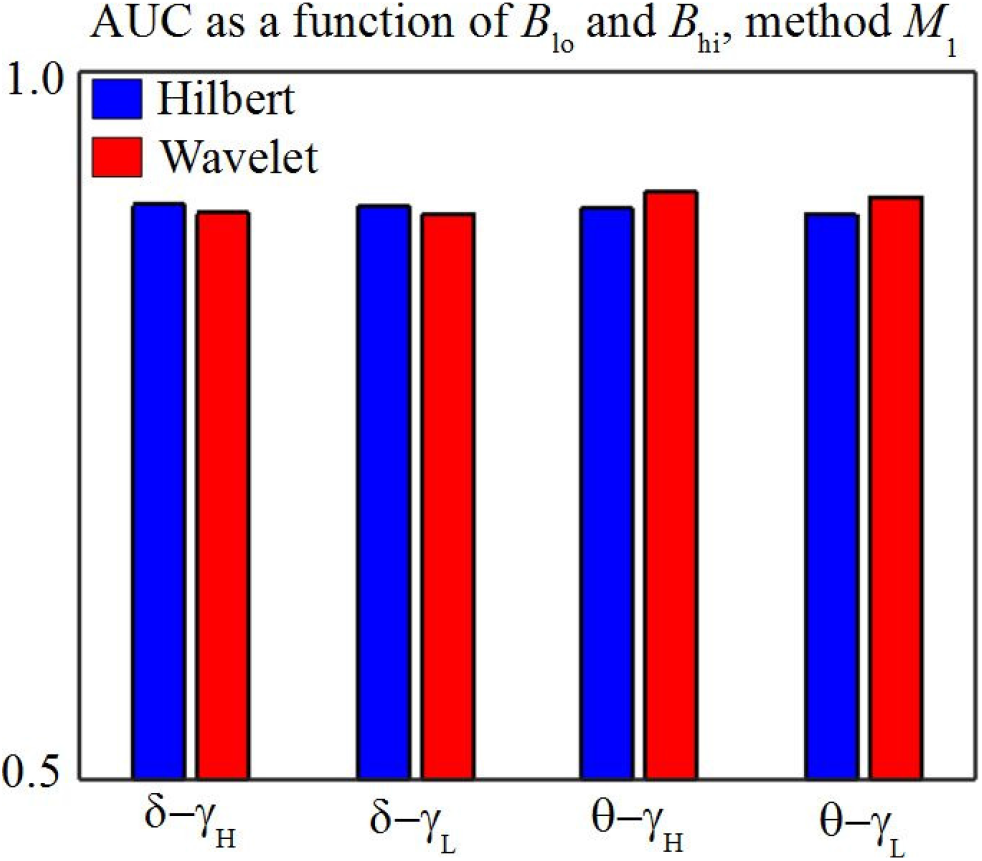
Areas under ROC curves for different values of the low- and high-frequency bands *B*_lo_ and *B*_hi_; blue bars are used for the Hilbert approach, and red bars for the wavelet approach. The values for the other parameters are: *χ* = 0.5, *T* =1s, *σ* =1, *f_s_* = 500 Hz, *f_I_* = 0 and PAC method *M*_1_.

**Figure 8:**
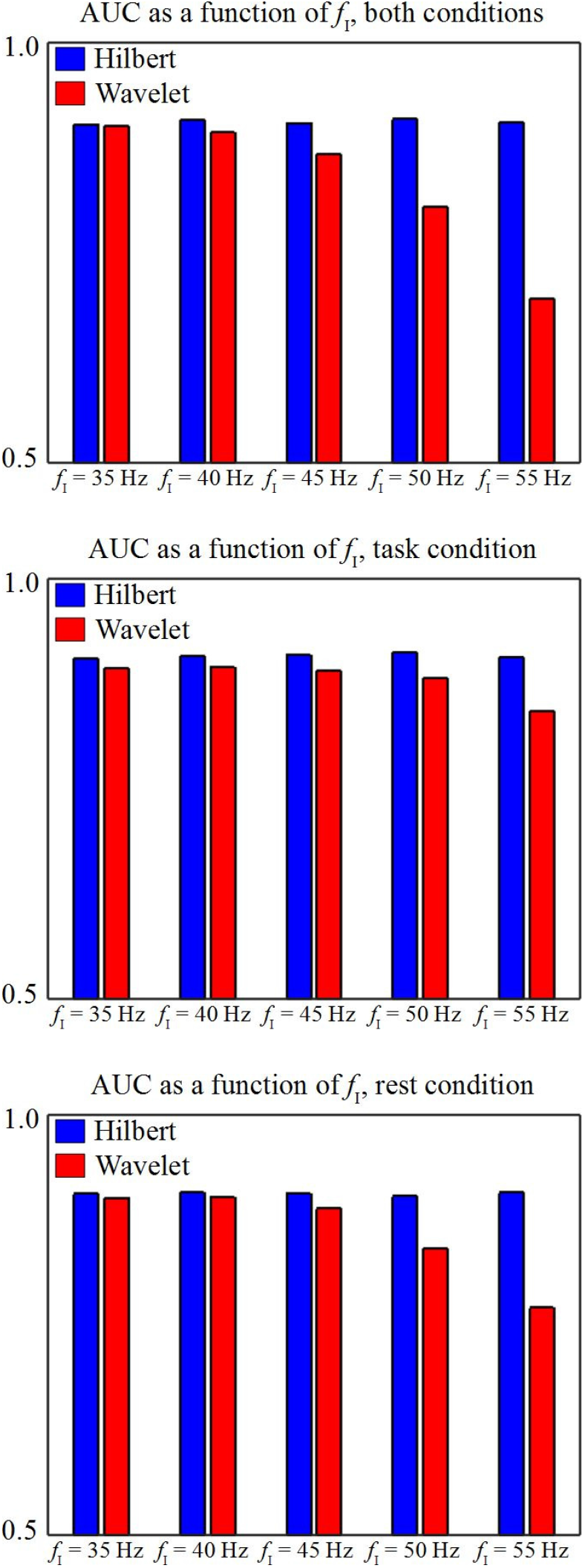
Areas under ROC curves for different values of frequency *f_I_* in Hz; blue bars are used for the Hilbert approach, and red bars for the wavelet approach. For the top plot, f/ = 0 for all realizations of *x*(*t*); for the middle plot, f = 0 only for the realizations of *x*(*t*) with *χ* < 1; for the bottom plot, f = 0 only for the realizations of *x*(*t*) with *χ* = 1. The values for the other parameters are: *χ* = 0.5, *σ* =1, *B*_lo_ = *δ*, *B*_hi_ = *γ*_*H*_, *T* = 1 s and *f_s_* = 500 Hz.

Given our simulation results, the factor mostly responsible for a disparity in detection performance between the two approaches is the presence of signal components in spectral regions near the high-frequency band under investigation. This finding, combined with the inherent wideband nature of electrophysiological measurements, provides a plausible explanation in real data analyses if there is a mismatch between significant brain activity found with each of the approaches. In the next subsection, we present the results of such an analysis.

### 3.3 MEG data

In figures 9, 10, 11 and 12 we present, for the Hilbert and the wavelet approaches, brain maps of multi-subject t-statistics that assess the difference between PAC during the VM condition and during rest, for four frequency band pairs: *δ*–*γ*_*H*_, *δ*–*γ*_*L*_, *θ*–*γ*_*H*_ and *θ*–*γ*_*L*_, respectively. We also indicate in figures 9 through 12 the brain regions where the use of each method resulted in a statistically significant effect. The most striking differences between the two approaches were observed with *δ*–*γ*_*H*_ and *δ*–*γ*_*L*_ coupling: in the former case (figure 9), the Hilbert approach enabled the detection of task-based changes in the cerebellum, around the visual cortex, in the right precentral sulcus, and near the left central sulcus (a decrease in the latter region, and an increase in all the others), while the wavelet approach yielded no regions with significant effects; as for the latter case (figure 10), we found with both approaches that in the vicinity of the central sulcus in both hemispheres there was a task-based decrease, but the spatial extent of the areas where there was this decrease was noticeably smaller with the wavelet approach (there was also an increase in PAC in small regions in the cerebellum and near the visual cortex that detected only with the Hilbert approach). As for the other two frequency band pairs, represented in figures 11 and 12, there were only very small active regions, such as a location near the left motor cortex with an increase in *θ*–*γ*_*H*_ PAC observed only with the wavelet approach, and another in the cerebellum with an increase in *θ*–*γ*_*L*_ PAC observed only with the Hilbert approach.

**Figure 9:**
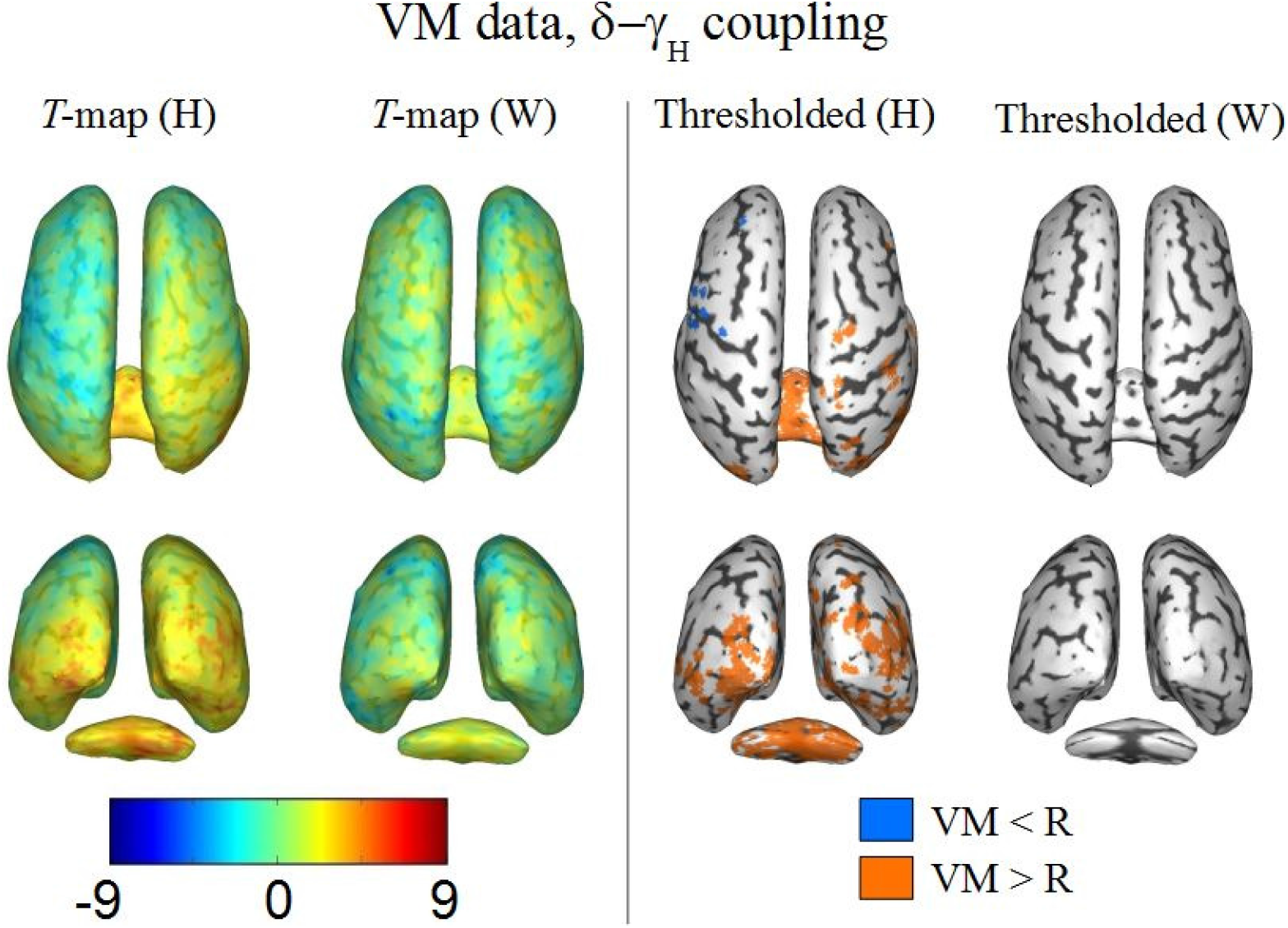
Multi-subject task-based changes in *δ*–*γ*_*H*_ coupling obtained from the VM data. *First column:* brain map of the group-level *t*-statistics computed with the Hilbert approach (H); *second column:* brain map of the group-level *t*-statistics computed with the wavelet approach (W); *third column:* brain locations with statistically significant task-based PAC changes computed with the Hilbert approach (H); *fourth column:* brain locations with statistically significant task-based PAC changes computed with the wavelet approach (W). For the last two columns, blue locations indicate task-based decrease in PAC, and orange locations represent task-based increase.

**Figure 10:**
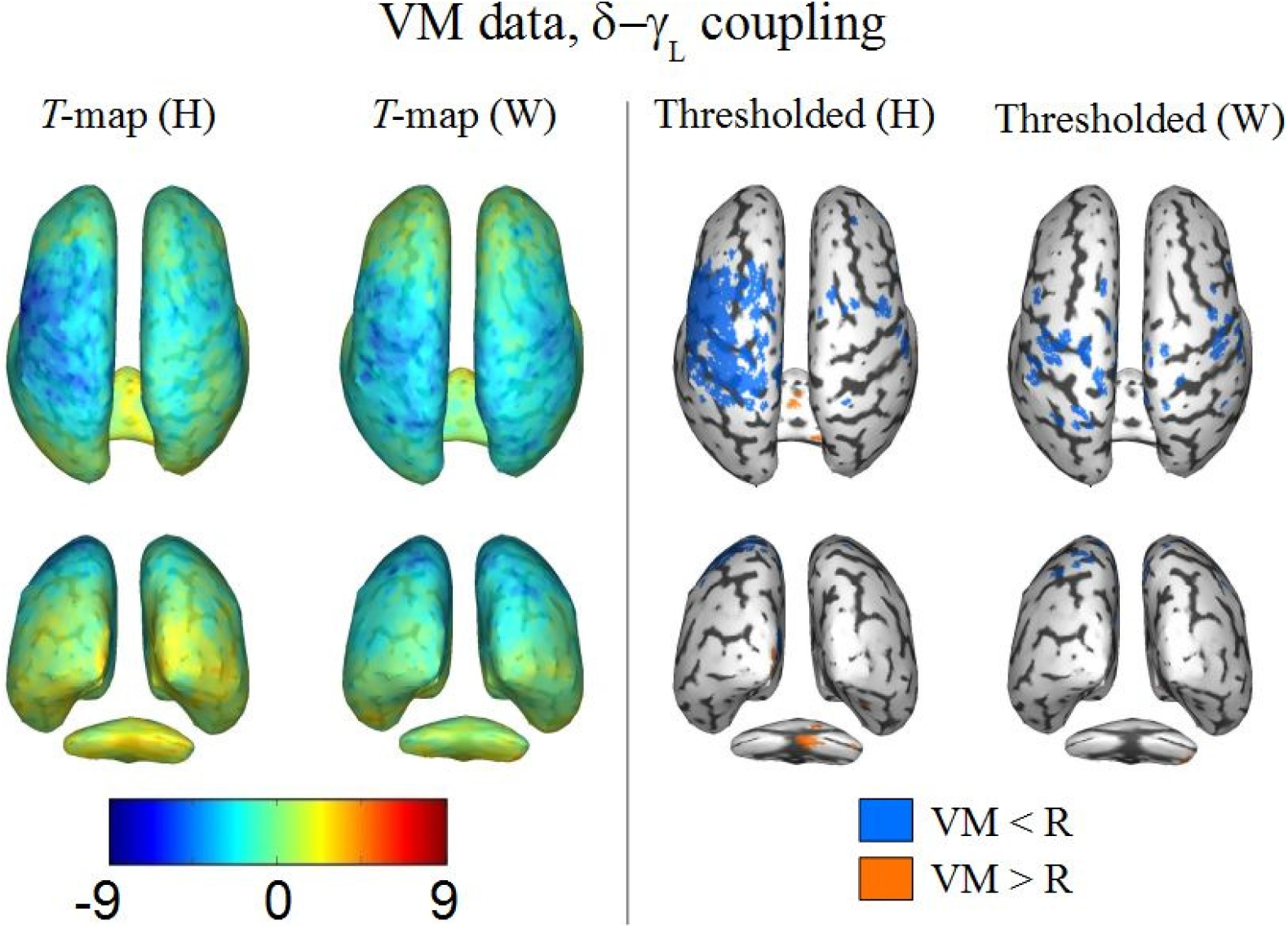
Similar to figure 9, for *δ*−*γ*_*L*_ coupling.

**Figure 11:**
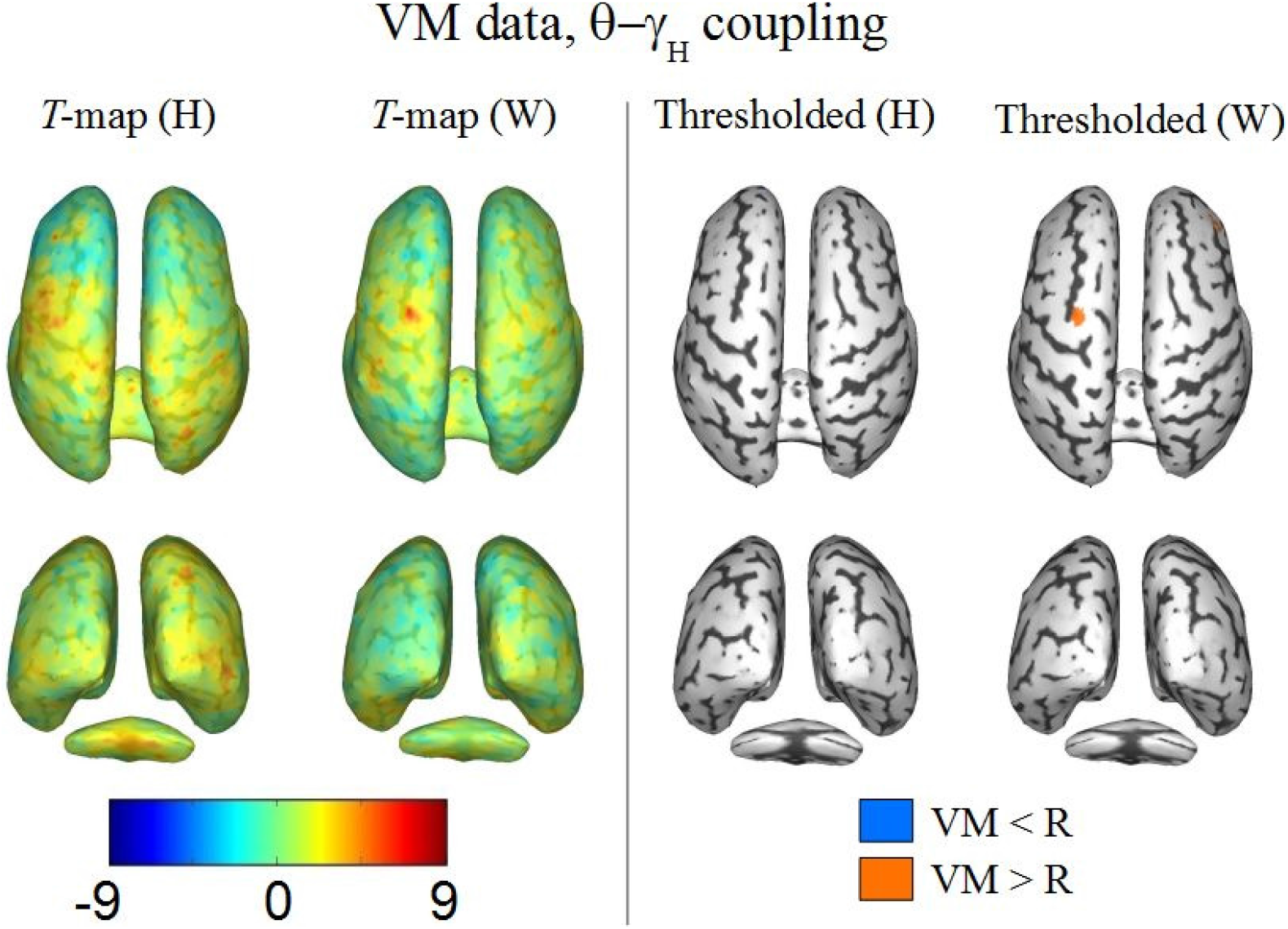
Similar to figure 9, for *θ*−*γ*_*H*_ coupling.

**Figure 12:**
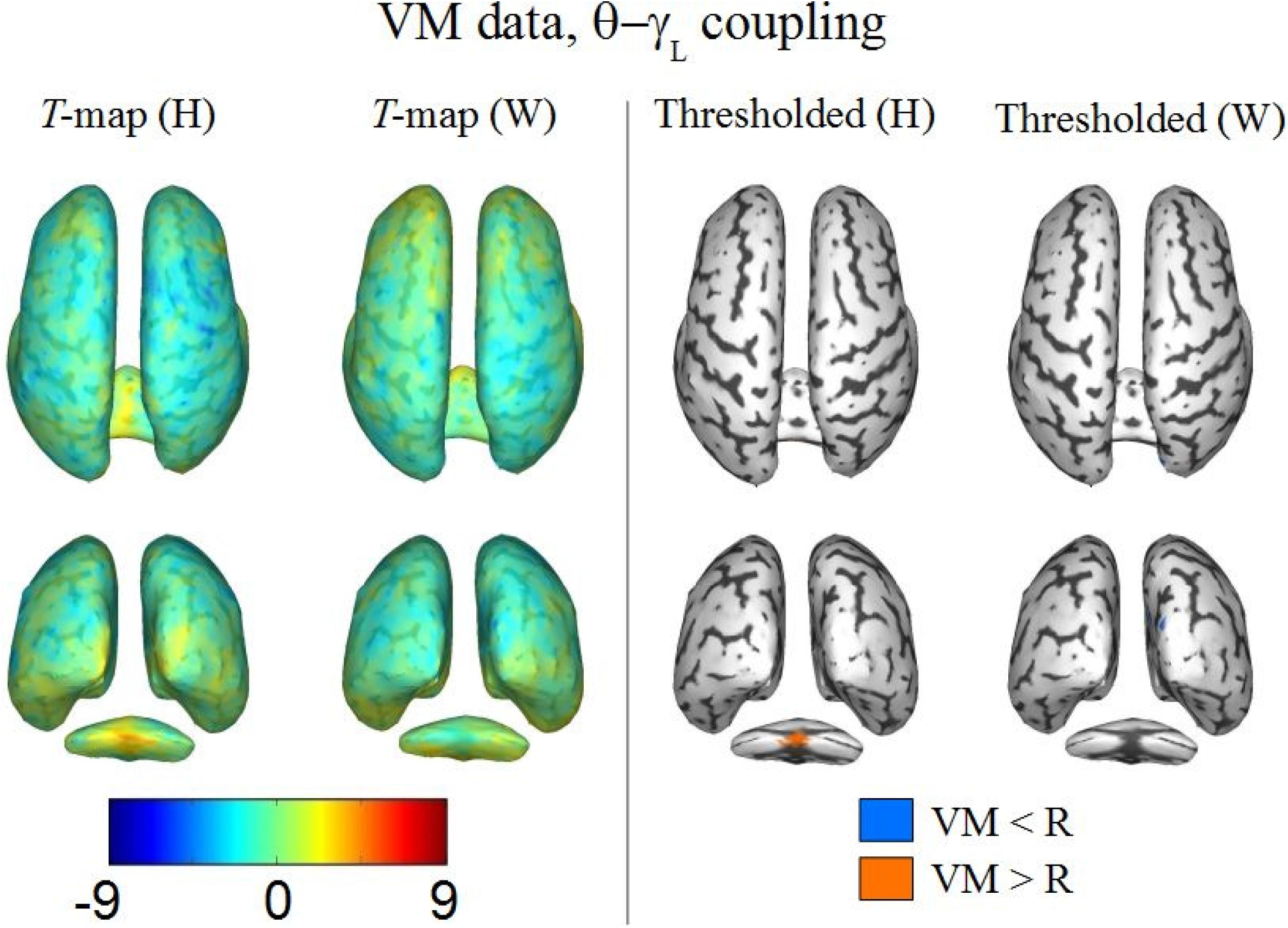
Similar to figure 9, for *θ*−*γ*_*L*_ coupling.

## 4 Discussion

Functional connectivity studies based on electrophysiological recordings, such as MEG and EEG, typically use as inputs to their analyses representations of brain activity signals within well-defined frequency bands. Two of the most commonly implemented approaches to extract these narrowband representations are band-pass filtering followed by computation of the Hilbert transform (the Hilbert approach, as we called it here) and convolution with Morlet wavelet kernels (the wavelet approach). In this work, our goal was to compare the Hilbert and the wavelet approaches in terms of their capabilities of detecting a specific form of interaction between brain signals – namely, phase-amplitude coupling, or PAC. In our results obtained with simulated brain activity, both methods yield comparable performance in a variety of practical scenarios; however, there was one feature of the synthetic time series that caused a noticeable degradation of the wavelet approach’s discrimination accuracy relative to that of the Hilbert approach – namely, the presence of a single sinusoidal wave oscillating at rates near the high-frequency band chosen for investigation, and uncorrelated with the active components within that band and the low-frequency one. This finding, combined with the inherent wideband nature of typical electrophysiology signals, is thus a potential explanation for the disparity we observed between the strongly coupled brain regions detected with each approach when they were applied to MEG recordings acquired during a visuomotor coordination experiment.

A common assumption in connectivity analyses performed with EEG or MEG data is that there is no preferred approach for obtaining the narrowband representations of time series, because the Hilbert and the wavelet approach can be expressed as linear convolutions of the signals of interest and are thus mathematically equivalent (Bruns, 2004; Kiebel et al., 2005). While the latter statement is correct, it does not correspond to how the approaches are actually implemented, and this can in fact result in fluctuations in performance depending on the approach selected, as our results demonstrate. From a theoretical perspective, although it is indeed possible to use Morlet wavelet kernels for band-pass filtering and for estimation of envelope amplitude and instantaneous phase (which would otherwise be computed with Hilbert transforms), these wavelets not only have frequency responses that deviate strongly from the properties usually desired in bandpass filters (namely, nearly flat passband, strong signal attenuation in the stopband, and very short transition bands), as figure 2 indicates, but they also perform only approximately the function of the Hilbert transform, which yields output signals with zero energy in negative frequencies – alternatives to the Morlet wavelet, such as the one proposed by Nakhnikian et al. (2016), are capable of dealing with the latter issue but not with the former. That bandpass filters designed to resemble Morlet wavelet kernels have indeed equivalent performance to the latter when estimating envelope amplitudes, as observed by (Bruns, 2004), should not come as a surprise, but neither should a noticeable disparity between the approaches in tasks other than amplitude estimation, or when filters with better frequency discrimination are employed.

Given our present knowledge, it is difficult to determine the full set of reasons why, for instance, both approaches perform well in phase-phase coupling analyses, as in (Quian Quiroga et al., 2002; Le Van Quyen et al., 2001), whereas for PAC computations the use of the Hilbert approach is clearly advantageous in some scenarios, as described in the present article. Our preliminary surveys (subsection 3.2) indicate a degradation in the performance of the wavelet approach when there are interfering signals only a bit slower than the high-frequency band, which is possibly linked to the frequency response properties of this approach (i.e. the transition band obtained with the Morlet wavelet is much longer than that provided by band-pass filtering, as displayed in figure 2), even though it remains to be determined the actual relationship between frequency response and poorer perfomance in PAC estimation. Phase-amplitude coupling compares two signals at different frequencies, whereas phase-phase coupling, as computed by Le Van Quyen et al. (2001), deals with narrowband signals with the same central frequency; keeping in mind that the most popular methods to evaluate phase coupling are usually based on phase differences (Tass et al., 1998; Lachaux et al., 1999), one might conjecture that the eventual leakage from frequencies outside the range of interest has comparable impact in both signals under analysis and it gets canceled out when computing this form of coupling. However, a much more extensive examination than the one carried out here will be necessary to determine, first, why the Hilbert approach performs better in PAC estimates when certain features of the brain signals are present, taking into account other possible factors for degradations in performance (phase and/or amplitude distortions, filter behavior near 0 Hz, and so on); this thorough examination may then be extended to other forms of coupling.

In our simulations, we created different configurations mainly by modifying the properties of the synthetic time series, such as noise levels, signal length, coupling strength, and sampling frequency. As for the parameters of both approaches, we either chose fixed values that were in line with what one commonly finds in the literature dealing with brain connectivity (e.g. filter length, type and window, with the Hilbert approach) or varied their values only in order to find optima to be used in the subsequent approach comparisons (specifically, number of wavelet kernels and their width). Regarding our procedure for this selection of parameters, one may raise objections that we believe are worth addressing: first, we acknowledge the possibility that the values we chose for the wavelet parameters are optimal exclusively for those simulation settings, therefore effects such as the poorer performance of the wavelet approach due to the interfering sinusoidal wave (figure 8) and the disparity of active regions between the approaches when applied to the VM data (figures 9-12) could be due merely to an inadequate choice of parameters. In that case, we must not exclude the possibility that a different choice of parameters, while benefiting the wavelet approach in some of the scenarios tested here (say, by reducing the disparity in the VM data maps), also result in a degradation in other simulation settings, and even in significant divergences between the approaches in other EEG/MEG acquisitions. A more specific issue is related to the value of the number of wavelet kernels *N*_ker_ = 1 we selected for all our analyses even though other values of the parameter provided higher areas under the ROC curves (table 2). Even taking into account considerations that more wavelet kernels would increase considerably processing times while providing only marginal improvements in accuracy, one might still argue that a higher *N*_ker_ could minimize the approach discrepancies, at least in theory. However, we find unlikely that convolving the time series with more wavelet kernels would, say, diminish the impact of the interfering sinusoids, especially given the frequency response properties of the Morlet wavelets, as illustrated in figure 2.

The differences in detection performance between the two approaches was also observed with MEG data obtained from a visuomotor coordination study. The most striking differences were found in the brain maps for PAC changes between *δ* and *γ*_*H*_ and between *δ* and *γ*_*L*_, where the wide areas that appeared to have strong effects with the Hilbert approach were either not found (figure 9) or had smaller spatial extent (10) with the wavelet approach. Given the findings we obtained with synthetic brain activity, especially the effects of the interfering sinusoids, one might feel tempted to assert that the images obtained with the Hilbert approach are the ones reflecting most accurately the true activity in the brain, but in order to validate such a statement more information should be gathered, e.g. with procedures that measure electrical brain activity with better spatial resolution, such as electrocorticography and depth electrodes. It would also be necessary to deal adequately with methodological issues that may prevent the correct identification of brain behavior with the available PAC methods (Aru et al., 2015), such as number of filters and their passband width, non-stationarity, and influence of signal power fluctuations in PAC modulations – in section 3.3, our goal with the VM data was essentially to observe the impact of the choice of approach with real MEG measurements. That being said, it should be pointed out that the some of the experimental outcomes of the present article coincide with other investigations dealing with the implications of visuomotor coordination in brain electrical activity: for instance, there is evidence of increased delta (Jerbi et al., 2007; Bradberry et al., 2009; Bourguignon et al., 2012; Mylonas et al., 2016) and gamma activity (Brovelli et al., 2017) in the motor cortex, and of gamma activity in the visual cortex (Barratt et al., 2017). Again, further analysis is necessary to better link our findings with those that appeared in the literature, particularly because the latter did not necessarily involve PAC computations.

As indicated in the preceding paragraphs, an interesting topic for further research, from a methodological point of view, is a more thorough examination of the properties of the Hilbert and the wavelet approaches, in order to discover the reasons for the variations in accuracy when computing PAC; eventually, the new information to be obtained can be applied to a comparison between these approaches to other forms of connectivity, such as phase synchrony, envelope amplitude correlations, and coherence. From a neuroscientific perspective, we intend to keep applying methods that estimate functional interactions to our visuomotor acquisitions, so that their impact on brain behavior may be better understood. Another possible topic worth exploring is the improvement of the current algorithms that extract narrowband representations of brain activity, in order to decrease processing times. As an example, a common technique to compute the Hilbert transform (as used by Matlab) is to evaluate the Fourier transform of the original signal, set to zero all components with negative frequencies, and then apply the inverse Fourier transform. Though accurate, this technique is inadequate in real-time situations, such as brain-computer interfaces, thus a faster alternative to it would be desirable.

